# Expansions of adaptive-like NK cells with a tissue-resident phenotype in human lung and blood

**DOI:** 10.1101/2019.12.20.883785

**Authors:** Nicole Marquardt, Marlena Scharenberg, Jeffrey E. Mold, Joanna Hård, Eliisa Kekäläinen, Marcus Buggert, Son Nguyen, Jennifer N. Wilson, Mamdoh Al-Ameri, Hans-Gustaf Ljunggren, Jakob Michaëlsson

## Abstract

Human adaptive-like “memory” CD56^dim^CD16^+^ NK cells in peripheral blood from cytomegalovirus-seropositive individuals have been extensively investigated in recent years and are currently explored as a treatment strategy for hematological cancers. However, treatment of solid tumors remains limited due to insufficient NK cell tumor infiltration, and it is unknown whether large expansions of adaptive-like NK cells that are equipped for tissue-residency and tumor-homing exist in peripheral tissues. Here, we show that human lung and blood contains adaptive-like CD56^bright^CD16^−^ NK cells with hallmarks of tissue-residency, including expression of CD49a. Expansions of adaptive-like lung tissue-resident (tr)NK cells were found to be present independently of adaptive-like CD56^dim^CD16^+^ NK cells and to be hyperresponsive towards target cells. Together, our data demonstrate that phenotypically, functionally, and developmentally distinct subsets of adaptive-like NK cells exist in human lung and blood. Given their tissue-related character and hyperresponsiveness, human lung adaptive-like trNK cells might represent a suitable alternative for therapies targeting solid tumors.

## Introduction

Natural killer (NK) cells are a crucial component of the innate immune system. They target and eliminate virus-infected and malignant cells, and boost immunity through the production of proinflammatory cytokines including IFN-γ and TNF. In recent years, the concept of adaptive-like or “memory” NK cells has emerged from studies in mice (1–4) and humans (5–10). These adaptive-like NK cells share a distinct phenotype and increased target cell responsiveness as well as having features of longevity and superior recall potential reminiscent of memory T cells (11).

Most studies of human adaptive-like NK cells have focused on subsets of NKG2C^+^(KIR^+^)CD56^dim^CD16^+^ NK cells, originally found to be expanded and stably imprinted in peripheral blood of approximately 30-40% of human CMV (HCMV) seropositive individuals (5,10). Adaptive-like NKG2C^+^(KIR^+^)CD56^dim^CD16^+^ NK cells in human peripheral blood have a distinctive phenotypic (5,10), epigenetic (8,9), and functional (8–10) profile compared to conventional NK cells and have been suggested to contribute to immunity against HCMV (1,12). Importantly, adaptive-like peripheral blood-derived CD56^dim^CD16^+^ NK cells (herein defined as adaptive-like CD56^dim^CD16^+^ pbNK cells) are currently explored for improving NK cell-mediated cancer therapies. While adoptive NK cell transfer showed optimistic results in the treatment of hematological malignancies (reviewed in (13)), targeting solid tumors was less successful due to poor migration to and infiltration into the tumor (reviewed in (13)). In these cases, adaptive-like NK cells with an increased capacity to infiltrate tissues e.g. through co-expression of tissue-specific ligands might be desirable. In fact, similar approaches for targeting solid tumors have recently been suggested for T_RM_ cells (14).

We and others recently identified a subset of tissue-resident CD49a^+^CD56^bright^CD16^−^ NK (trNK) cells in the human lung (15,16). The human lung is a frequent site of acute infections, including infections with viruses such as influenza virus, respiratory syncytial virus (RSV), and HCMV, as well as serving as a reservoir for latent HCMV infection (17). Although human CD56^dim^CD16^+^ lung NK cells are hyporesponsive to *ex vivo* target cell stimulation (18), exposure of human lung NK cells to virus-infected cells is likely to result in functional NK cell priming and expansion of distinct NK cell subsets. Indeed, increased polyfunctional responses have been observed in CD16^−^ lung NK cells following *in vitro* infection with IAV (16,19). However, the presence of expansions of functional adaptive-like trNK cells in the human lung is to date unknown.

Here, we identify and examine a CD49a^+^KIR^+^NKG2C^+^CD56^bright^CD16^−^ NK cell population with features of tissue-resident NK cells in human lung and blood, which is distinct from adaptive-like CD56^dim^CD16^+^ pbNK cells. In donors with expansions of adaptive-like CD49a^+^ lung trNK cells, small but detectable frequencies of adaptive-like CD49a^+^ NK cells were observed in paired peripheral blood. While adaptive-like CD56^dim^CD16^+^ pbNK cells (as commonly identified in peripheral blood of HCMV-seropositive donors) and adaptive-like CD49a^+^ trNK cells in lung and blood shared a common core gene signature, we identified several unique features of each population indicating that they may represent developmentally distinct populations. Notably, NK cells from donors with an adaptive-like trNK cell expansion in the lung were hyperresponsive towards target cells. Thus, we provide evidence indicating that CD49a^+^KIR^+^NKG2C^+^CD56^bright^CD16^−^ trNK cells in the human lung represent a distinct population of adaptive-like NK cells with potential implications in lung surveillance and future treatment options of solid tumors.

## Results

### Adaptive-like NK cells can be identified in human lung

We first set out to investigate whether expansions of adaptive-like KIR^+^NKG2C^+^ NK cells could be identified in the human lung. The majority of NK cells in the lung are phenotypically similar to pbNK cells (CD69^−^CD56^dim^CD16^+^), suggesting that these cells may recirculate between the lungs and peripheral blood (18). Accordingly, circulating populations of expanded adaptive-like CD56^dim^CD16^+^ NK cells could also be identified in both the peripheral blood and lungs from patients undergoing surgery for suspected lung cancer (Fig. 1A). Surprisingly, KIR and NKG2C were also co-expressed on less differentiated CD56^bright^CD16^−^ lung NK cells in many of the patients included in the study, with varying frequencies between donors (Fig. 1B, C) (see Supplementary Fig. 1A for the gating strategy to identify KIR^+^NKG2C^+^ CD56^bright^CD16^−^ and CD56^dim^CD16^+^ NK cells). In several donors the frequencies of KIR^+^NKG2C^+^CD56^bright^CD16^−^ NK cells in human lung vastly exceeded those previously described for CD16^−^ NK cells in the liver (7), with up to 98% of CD56^bright^CD16^−^ lung NK cells co-expressing KIR and NKG2C (Fig. 1 A, B).

**Figure 1:**
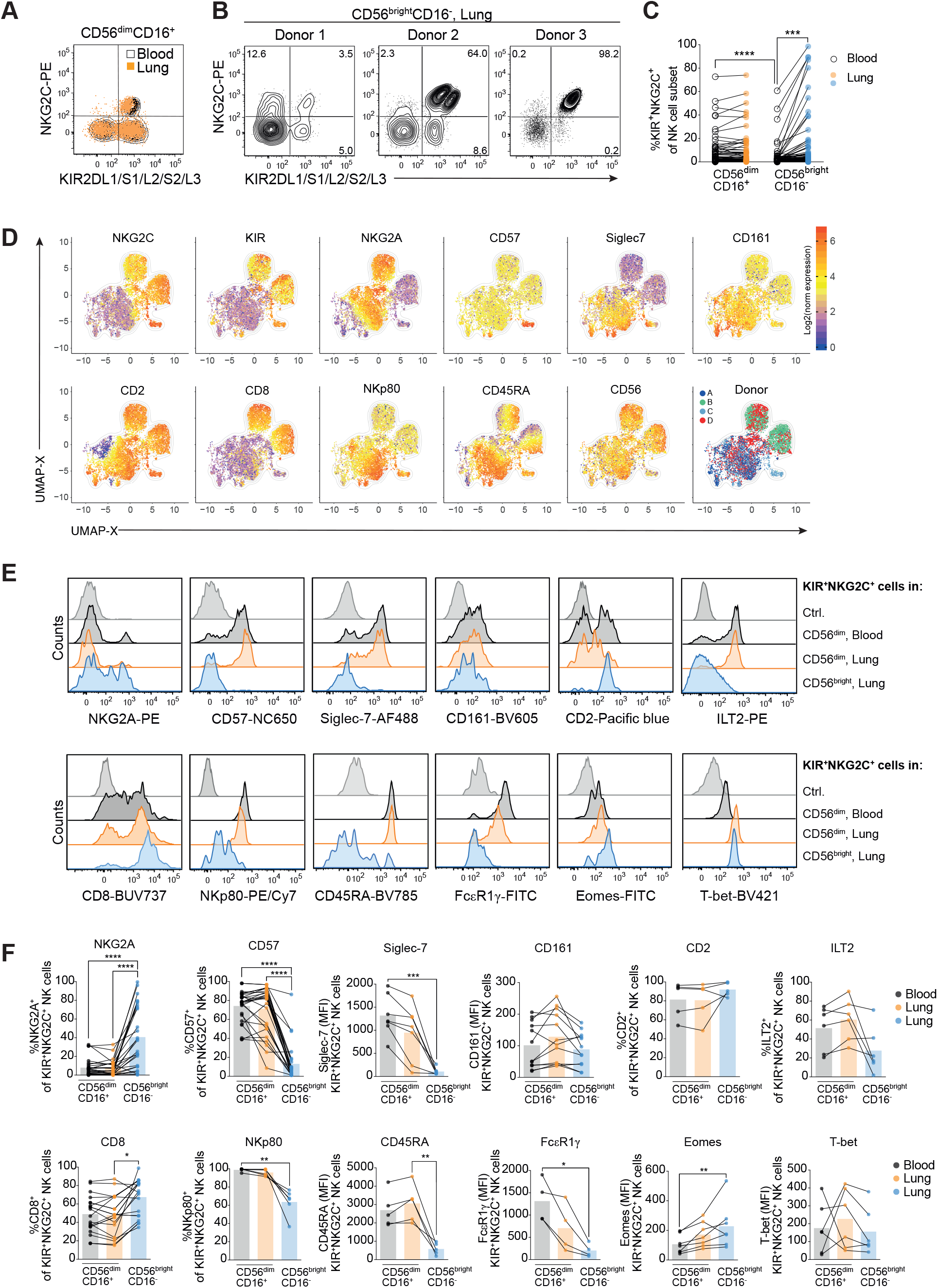
Adaptive-like KIR^+^NKG2C^+^ NK cells exist in the CD56^bright^CD16^−^ NK cell subset in the human lung. **(A)** Representative overlay displaying pan-KIR and NKG2C expression on CD56^dim^CD16^+^ NK cells in paired blood (black contour) and lung (orange). **(B)** Representative dot plots displaying pan-KIR and NKG2C expression on CD56^bright^CD16^−^ NK cells in the lungs of three different donors. **(C)** Summary of data showing the frequencies of KIR^+^NKG2C^+^ NK cells in CD56^dim^CD16^+^ and CD56^bright^CD16^−^ NK cells in paired blood and lung (n = 77). Friedman test, Dunn’s multiple comparisons test. ***p<0.001, ****p<0.0001. **(D)** UMAP analysis of CD56^bright^CD16^−^ lung NK cells from four donors with 2,000 events/donor (942 events in one of the donors). UMAPs were constructed using expression of Siglec-7, CD8, CD2, CD57, CD161, NKG2C, CD56, CD45RA, NKG2A and NKp80. Color scale shows log2(normalized expression + 1). **(E)** Representative histograms and **(F)** summary of data showing surface expression of NKG2A (n = 27), CD57 (n = 27), Siglec-7 (n = 7), CD161 (n = 12), CD2 (n = 5), ILT2 (n = 6), CD8 (n = 20), NKp80 (n = 6), CD45RA (n = 5), and intracellular expression of FcεR1γ (n = 4), Eomes (n = 7) and T-bet (n = 6) in KIR^+^NKG2C^+^ NK cells in CD56^dim^CD16^+^ blood (grey) and lung (orange) NK cells and CD56^bright^CD16^−^ lung NK cells (blue). Friedman test, Dunn’s multiple comparisons test. *p<0.05, **p<0.01, ***p<0.001, ****p<0.0001.

Next, we assessed phenotypic features of KIR^+^NKG2C^+^CD56^bright^CD16^−^ lung NK cells in an unbiased manner using high dimensional flow cytometry. Uniform manifold approximation and expression (UMAP) analysis revealed a distinct subset of cells with an expression pattern consistent with adaptive-like NK cells found in peripheral blood and liver, including low expression of Siglec-7 and CD161, and high expression of NKG2C, KIRs, and CD2 (7,8,20,21) (Fig. 1D). Unlike adaptive-like CD56^dim^CD16^+^ pbNK cells, KIR^+^NKG2C^+^CD56^bright^CD16^−^ NK cells expressed lower levels of CD45RA and NKp80, and higher levels of CD8 (Fig. 1D). In addition, to these expression patterns, manual analysis of individual samples additionally confirmed low expression of ILT2 and FcεR1γ, as compared to KIR^+^NKG2C^+^CD56^dim^CD16^+^ lung NK cells (Fig. 1E, F). Furthermore, KIR^+^NKG2C^+^CD56^bright^CD16^−^ lung NK cells displayed low expression of ILT2 and FcεR1γ and high expression of Eomes and NKG2A, but similar levels in T-bet expression, revealing a phenotype distinct from human adaptive-like KIR^+^NKG2C^+^CD16^−^ NK cells in the liver (7) (Fig. 1E, F). Together, our data reveal the presence of a previously uncharacterized population of an adaptive-like NK cell subset, herein identified as KIR^+^NKG2C^+^CD56^bright^CD16^−^, in the human lung, which is distinct from adaptive-like CD56^dim^CD16^+^ pbNK cells.

### Adaptive-like CD56^bright^CD16^−^ NK cells in human lung are tissue-resident

TrNK cells in human lung are characterized by expression of CD69 and the integrins CD49a and CD103 (15,16), while adaptive-like NK cells have not yet been described within this subset. Strikingly, the vast majority of the distinct population of NKG2C^+^CD56^bright^CD16^−^ NK cells identified by UMAP analysis co-expressed CD69 (80%) and CD49a (77%), and a substantial proportion (38%) also co-expressed CD103 (Fig. 2A, B), suggesting tissue-residency of adaptive-like CD56^bright^CD16^−^ NK cells in the lung. KIR^+^NKG2C^+^ NK cells co-expressing CD49a, CD69, or CD103, were mainly CD56^bright^CD16^−^ (Fig. 2C, D), further demonstrating that they are clearly distinct from adaptive-like CD56^dim^CD16^+^ pbNK cells.

**Figure 2:**
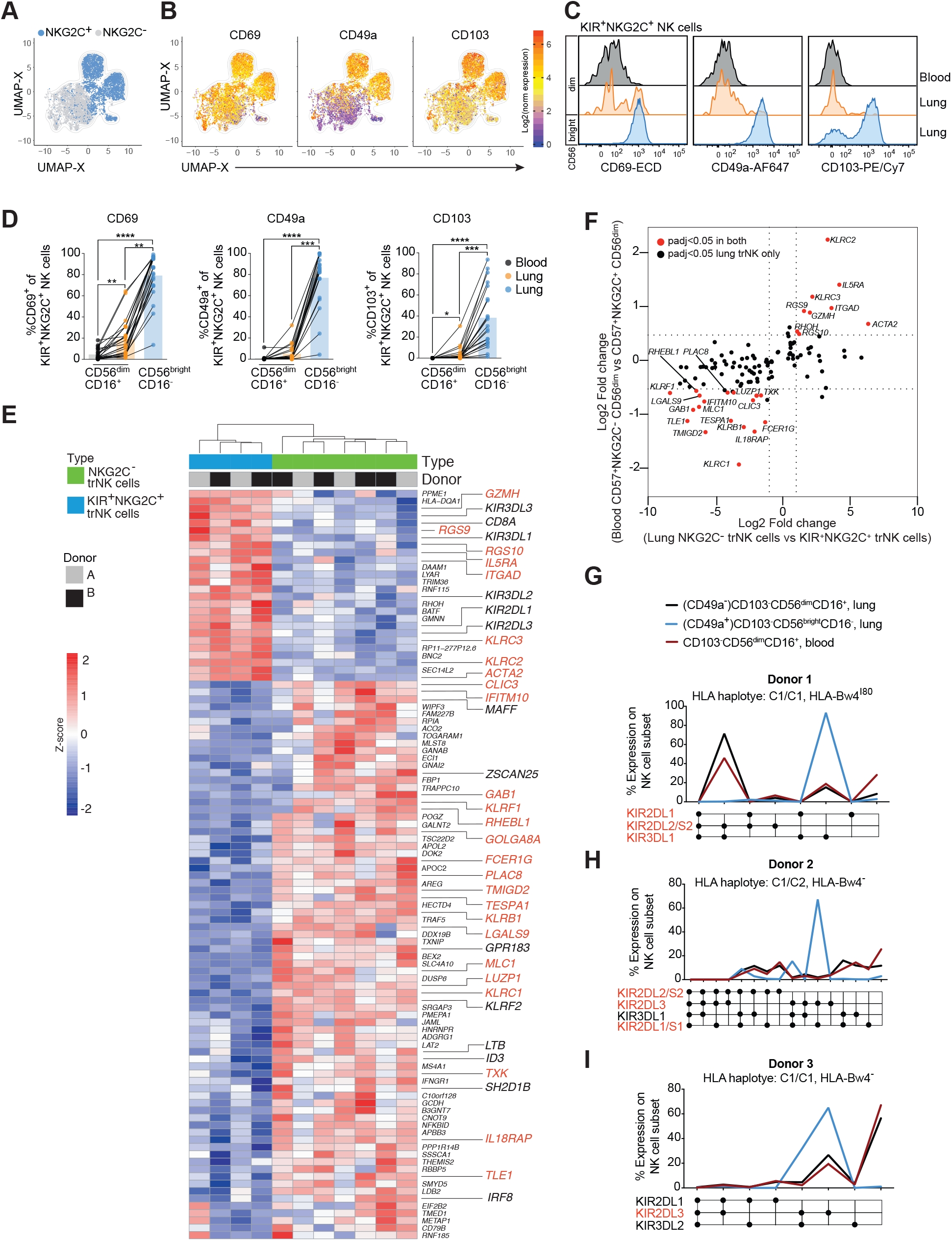
Adaptive-like CD56^bright^CD16^−^ lung NK cells are tissue-resident. **(A)** Overlay of UMAPs from data from Figure 1, showing the position of NKG2C^+^ (blue) and NKG2C^−^ (grey) populations among CD56^bright^CD16^−^ NK cells. **(B)** Expression of the tissue-residency markers CD69, CD49a, and CD103 within the UMAP of CD56^bright^CD16^−^ lung NK cells. **(C)** Representative histograms and **(D)** summary of data showing the expression of the tissue-residency markers CD69 (n = 23), CD49a (n=21) and CD103 (n = 21) on CD56^dim^CD16^+^ blood (grey) and lung (orange) NK cells and CD56^bright^CD16^−^ lung NK cells (blue), respectively. Friedman test, Dunn’s multiple comparisons test. *p<0.05, **p<0.01, ***p<0.001, ****p<0.0001. **(E)** Heatmap showing 102 differentially expressed genes between KIR^+^NKG2C^+^ trNK cells and KIR^−^NKG2C^−^ trNK cells in the human lung. Differentially expressed genes shared with CD57^+^NKG2C^+^CD56^dim^CD16^+^ adaptive-like NK cells in blood (from GSE117614) are highlighted in red. **(F)** Log2 fold-change for non-adaptive (NKG2C^−^) trNK cells vs adaptive-like trNK cells in lung against log2 fold change for CD57^+^NKG2C-vs adaptive-like CD57^+^NKG2C^+^ CD56^dim^ NK cells in blood. Data for CD56^dim^ NK cells in peripheral blood are from GSE117614 (25). **(G-I)** Single KIR expression analysis on CD56^dim^CD16^+^ NK cells from peripheral blood (red), CD49a^−^CD103^−^CD56^dim^CD16^+^ or CD103^−^CD56^dim^CD16^+^ (black) and CD49a^+^CD103^+^CD56^bright^CD16^−^ or CD103^+^CD56^bright^CD16^−^ (blue) NK cells in matched lung of three different donors. Educating KIR are highlighted in red. **(G)** Donor with expansions of self-KIR^+^ NK cells both in the CD56^dim^CD16^+^ subset in paired blood and lung and in the CD56^bright^CD16^−^ NK cell subset in the lung. **(H)** Donor with an expansion of self-KIR^+^ NK cells exclusively in the CD56^bright^CD16^−^ NK cell subset in the lung. **(I)** Donor with expansions of self-KIR^+^ NK cells both in the CD56^dim^CD16^+^ and CD56^bright^CD16^−^ subsets in paired blood and lung, respectively.

To further characterize adaptive-like trNK cells in the lung, we compared the gene expression profiles of sorted adaptive KIR^+^NKG2C^+^ and non-adaptive KIR^−^ NKG2C^−^ trNK cells in human lung using RNA sequencing (Fig. 2E, F; see sorting strategy in Supplementary figure 1B, C). 102 genes were differentially expressed (p<0.05, log2FC>1), including several *KIR* genes, *CD8A*, *GPR183* (EBI2)*, IRF8* and *SH2D1B* (EAT2), and genes encoding for the transcription factors MafF (*MAFF*) and ZNF498 (*ZSCAN25*). GPR183 has been demonstrated to be crucial for tissue-specific migration of innate lymphoid cells (22), while EAT2 expression has previously been shown to be downregulated in adaptive-like CD56^dim^CD16^+^ pbNK cells (8). Interestingly, while upregulation of both *IRF8* and *THEMIS2* has been reported to be essential for NK cell-mediated responses against MCMV infection (23,24), gene expression of both molecules was low in adaptive-like trNK cells in human lung (Fig. 2E), indicating different activation pathways in adaptive-like NK cells in mice and humans. Furthermore, we compared differentially expressed genes in adaptive-like lung trNK cells *versus* CD56^dim^CD16^+^ pbNK cells and observed an overlap of approximately one third of the differentially expressed genes (GSE117614) (25). These genes included *KLRC2* (NKG2C), *KLRC3* (NKG2E), *IL5RA*, *GZMH*, *ITGAD* (CD11d), *RGS9*, *RGS10* (upregulated), and *KLRB1* (CD161), *KLRC1* (NKG2A), *KLRF1* (NKp80), *TMIGD2*, *IL18RAP*, *FCER1G*, *MLC1*, *CLIC3* and *TLE1* (downregulated) (Fig. 2F). These results demonstrate that adaptive-like lung trNK and CD56^dim^CD16^+^ pbNK cells share a common core gene set specific for adaptive-like NK cells.

Adaptive-like NK cells in peripheral blood and in the human liver commonly have a distinct KIR expression profile which is dominated by KIRs that bind to self-HLA class I (self-KIRs) (7,10,26). In the lung, analysis of single KIR expression on CD16^−^ and CD16^+^ NK cell subsets in donors with high frequencies of adaptive-like lung trNK cells revealed that the former subset displayed unique KIR expression patterns compared to CD16^+^ NK cells in paired blood or lung (Fig. 2G-I; Supplementary Fig. 1D for the gating strategy). High expression of KIRs on the adaptive-like trNK cells was limited to self-KIR, identical to the phenotype described for adaptive-like NK cells in liver and peripheral blood. Notably, even in a donor with adaptive-like NK cell expansions of both trNK cells and pbNK cells (Fig. 2G), the KIR-expression profile differed between the two subsets. These results suggest a subset-specific development and/or differentiation of adaptive-like NK cells in blood and lung.

Together, CD49a^+^KIR^+^NKG2C^+^CD56^bright^CD16^−^ trNK cells in the human lung exhibit a unique profile of activating and inhibitory NK cell receptors (e.g. NKG2A, KIR, NKp80), as well as adaptor, signaling, and effector molecules (FcεR1γ, SH2D1B, granzyme H). This indicates that these are bona fide adaptive-like trNK cells which are distinct from adaptive-like CD56^dim^CD16^+^ pbNK cells, demonstrating the presence of an as of yet unexplored NK cell population in the human lung.

### Lung adaptive-like trNK cells are hyperresponsive to target cells

In order to determine whether adaptive-like lung trNK cells differed functionally from non-adaptive CD56^bright^CD16^−^ lung trNK cells, we first compared expression levels of genes associated with functional competence (Fig. 3A). Gene expression in adaptive-like lung trNK cells was higher for *IFNG*, *IL32*, *XCL1* and *GZMH* (granzyme H), and lower for *GNLY* (granulysin), *GZMA* (granzyme A), *GZMK* (granzyme K), *IL2RB* (CD122) and *IL18RAP* as compared to non-adaptive lung trNK cells (Fig. 3A). These results suggest a potential cytokine-mediated priming of adaptive-like trNK cells *in vivo*, e.g. by IL-12 and IL-18, which are potent inducers of IFN-γ and IL-32 in NK cells (27). While expression levels of the effector molecules *PRF1* (perforin) and *GZMB* (granzyme B) did not differ between adaptive- and non-adaptive-like CD56^bright^CD16^−^ NK cells at transcriptome level (Fig. 3A), *ex vivo* protein expression was increased for granzyme B in adaptive-like trNK cells (Fig. 3B, C). This indicated a potential cytotoxic function in this particular subset, despite lung NK cells generally being characterized as hyporesponsive to target cell stimulation (16,18). Intriguingly, NK cells from donors with an expansion of adaptive-like trNK cells in the lung degranulated stronger and produced more TNF compared to those from donors without such expansions (Fig. 3D). In particular NK cells co-expressing CD49a and KIR degranulated strongly and produced high levels of TNF upon target cell stimulation (Fig. 3D-F). This hyperresponsiveness of adaptive-like lung trNK cells was independent from co-expression of CD103, since similar activation levels were observed between CD49a^+^CD103^−^ and CD49a^+^CD103^+^ KIR^+^NKG2C^+^ NK cells (Fig. 3F-H). Furthermore, blood NK cells from donors with expanded adaptive-like lung trNK cells also responded stronger to target cells as compared to donors without such expansions (Fig. 3F-H). Taken together, these results revealed that the presence of expanded adaptive-like trNK cells in the lung is linked to hyperresponsivity towards target cells, implicating a role of these cells in active immune regulation within the lung.

**Figure 3:**
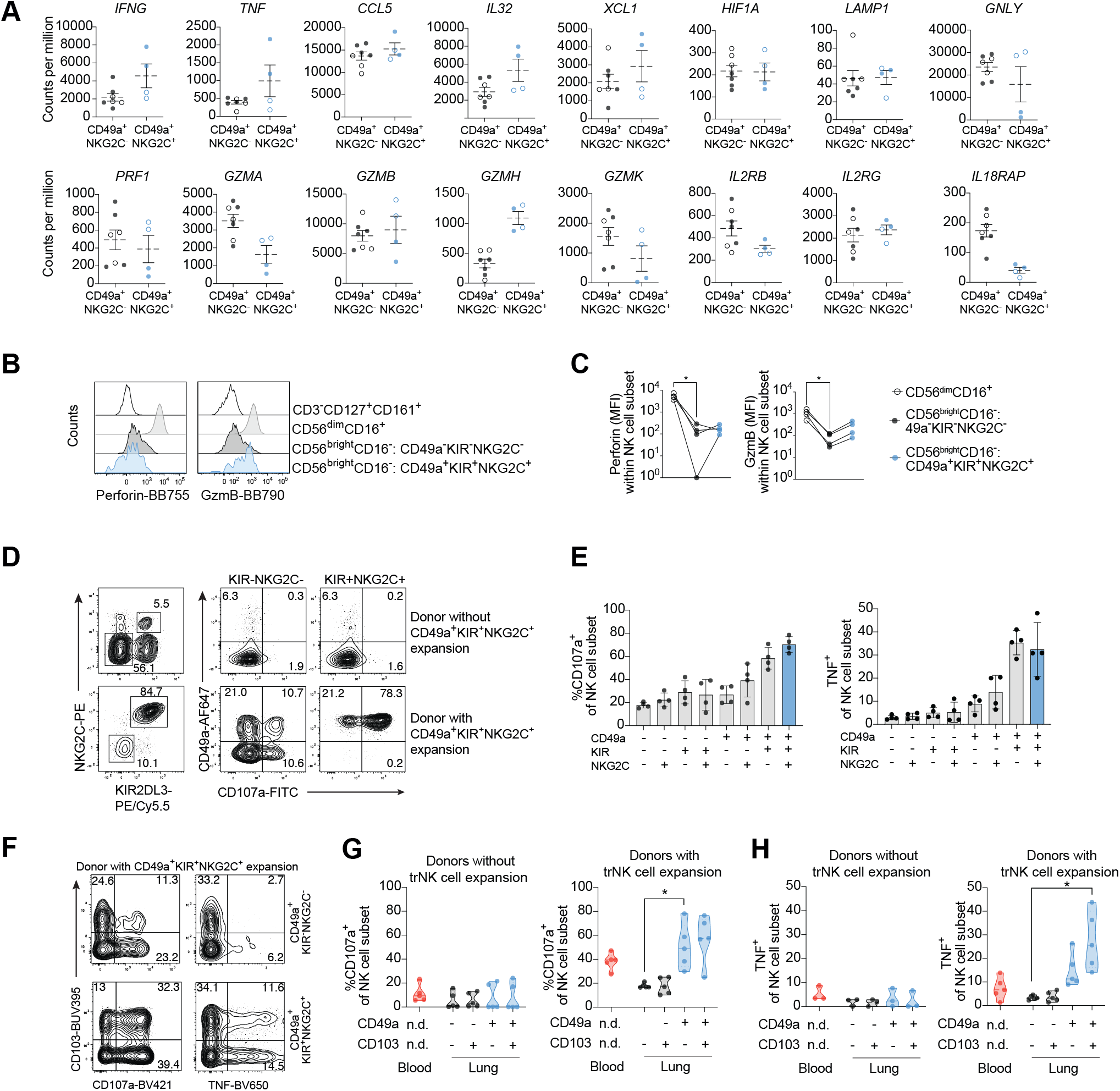
Adaptive-like lung trNK cells are highly functional. **(A)** Gene expression levels (counts per million reads) for selected genes associated with functional capacity are shown for non-adaptive CD49a^+^KIR^−^NKG2C^−^ and adaptive-like CD49a^+^KIR^+^NKG2C^+^ lung trNK cells (clear circles: CD49a^+^CD103^−^ NK cells, filled circles: CD49a^+^CD103^+^ NK cells). Mean ± SEM is shown. **(B)** Representative histograms and **(C)** summary of data displaying expression of perforin and granzyme B (GzmB) (n = 4) in CD56^dim^CD16^+^ and in non-adaptive CD49a^−^CD56^bright^CD16^−^ as well as adaptive-like CD49a^+^CD56^bright^CD16^−^ NK cells, respectively, in human lung *ex vivo*. CD14^−^CD19^−^CD3^−^CD45^+^CD127^+^CD161^+^ cells were gated as controls in (B). Friedman test, Dunn’s multiple comparisons test. *p<0.05. **(D)** Representative dot plots showing expression of CD107a and CD49a on KIR^−^NKG2C^−^ and KIR^+^NKG2C^+^ NK cells in a donor without (upper panel) and with (lower panel) expansion of adaptive-like trNK cells in the lung (expression KIR and NKG2C are displayed in the left panel for each of the two donors). **(E)** Summary of data showing the frequency of K562 target cell-induced CD107a^+^ (left) and TNF^+^ (right) NK cell subsets from donors with NK cell expansions in the human lung. Responses by unstimulated controls were subtracted from stimulated cells (n = 4). Mean ± SD is shown. **(F)** Representative dot plots showing expression of CD107a and TNF vs CD103 on non-adaptive CD49a^+^KIR^−^ NKG2C^−^ (upper panel) or adaptive-like CD49a^+^KIR^+^NKG2C^+^ (lower panel) bulk NK cells in a donor with an expansion of adaptive-like trNK cells in the lung. **(G, H)** Summary of data showing the frequencies of **(G)** CD107a^+^ and **(H)** TNF^+^ NK cells in blood NK cells and in subsets of lung NK cells (CD49a^−^CD103^−^, expressing either CD49a or CD103, or CD49a^+^CD103^+^) from donors without (left panels, n = 5 for CD107a, n= for TNF) or with (right panels, n = 5) expansions of KIR^+^NKG2C^+^ trNK cells in the lung. Responses by unstimulated controls were subtracted from stimulated cells. (G, H) Violin plots with quartiles and median are shown. Friedman test, Dunn’s multiple comparisons test. *p<0.05.

### Adaptive-like CD49a^+^CD56^bright^CD16^−^ NK cells can be identified in matched patient peripheral blood

As a hallmark of tissue-resident cells, CD49a is commonly expressed on subsets of T cells and NK cells in non-lymphoid compartments such as the lung (15,16,19,28), liver (7), skin (29), uterus (30), and intestine (31), but not in peripheral blood. Intriguingly, however, we identified a small subset of CD49a^+^KIR^+^NKG2C^+^ NK cells within the CD16^−^ NK cell population in paired peripheral blood of donors harboring expansions of adaptive-like trNK cells in the lung (Fig. 4A, B, see gating strategy in Supplementary Fig. 1E). The frequencies of CD49a^+^KIR^+^NKG2C^+^CD16^−^ NK cells in peripheral blood (herein identified as CD49a^+^ pbNK cells) were overall considerably lower as compared to either adaptive-like lung trNK cells or CD56^dim^CD16^+^ pbNK cells, respectively (Fig. 4B). We observed that 18.6% and 25.6% of all donors had an expansion (identified as outliers) of adaptive-like CD49a^+^CD56^bright^CD16^−^ NK cells in lung and paired peripheral blood, respectively. In comparison, 20.9% and 15.1% of all donors had an expansion of adaptive-like CD56^dim^CD16^+^ pbNK cells in lung and paired blood, respectively (Fig. 4B). Interestingly, expansions of adaptive-like CD49a^+^CD56^bright^CD16^−^ NK and CD56^dim^CD16^+^ pbNK cells were virtually mutually exclusive in donors (Fig. 4C). However, there was a substantial overlap within each of these subsets between lung and peripheral blood (Fig. 4C), that is, e.g. a high likelihood of having an expansion of adaptive-like trNK cells in the lung if an expansion of adaptive-like CD49a^+^ NK cells was present in the paired blood, and *vice versa*. This distinct distribution of adaptive-like NK cell populations per donor suggests adaptive-like CD49a^+^CD56^bright^CD16^−^ and CD56^dim^CD16^−^ pbNK cells are developing independently from each other. We next analyzed the phenotype of adaptive-like CD49a^+^CD56^bright^CD16^−^ pbNK cells and found intermediate expression of CD57 with relatively low co-expression of NKG2A (Fig. 4D, E), in contrast to lower expression of CD57 and higher expression of NKG2A in the adaptive-like lung trNK cells (Fig. 1E, F). Furthermore, adaptive-like CD49a^+^CD56^bright^CD16^−^ pbNK cells differed from their counterpart in lung by low expression of both CD69 and CD103 (Fig. 4E, D), consistent with known phenotypic differences between tissue-resident and circulatory lymphocyte populations (15,32–34).

**Figure 4:**
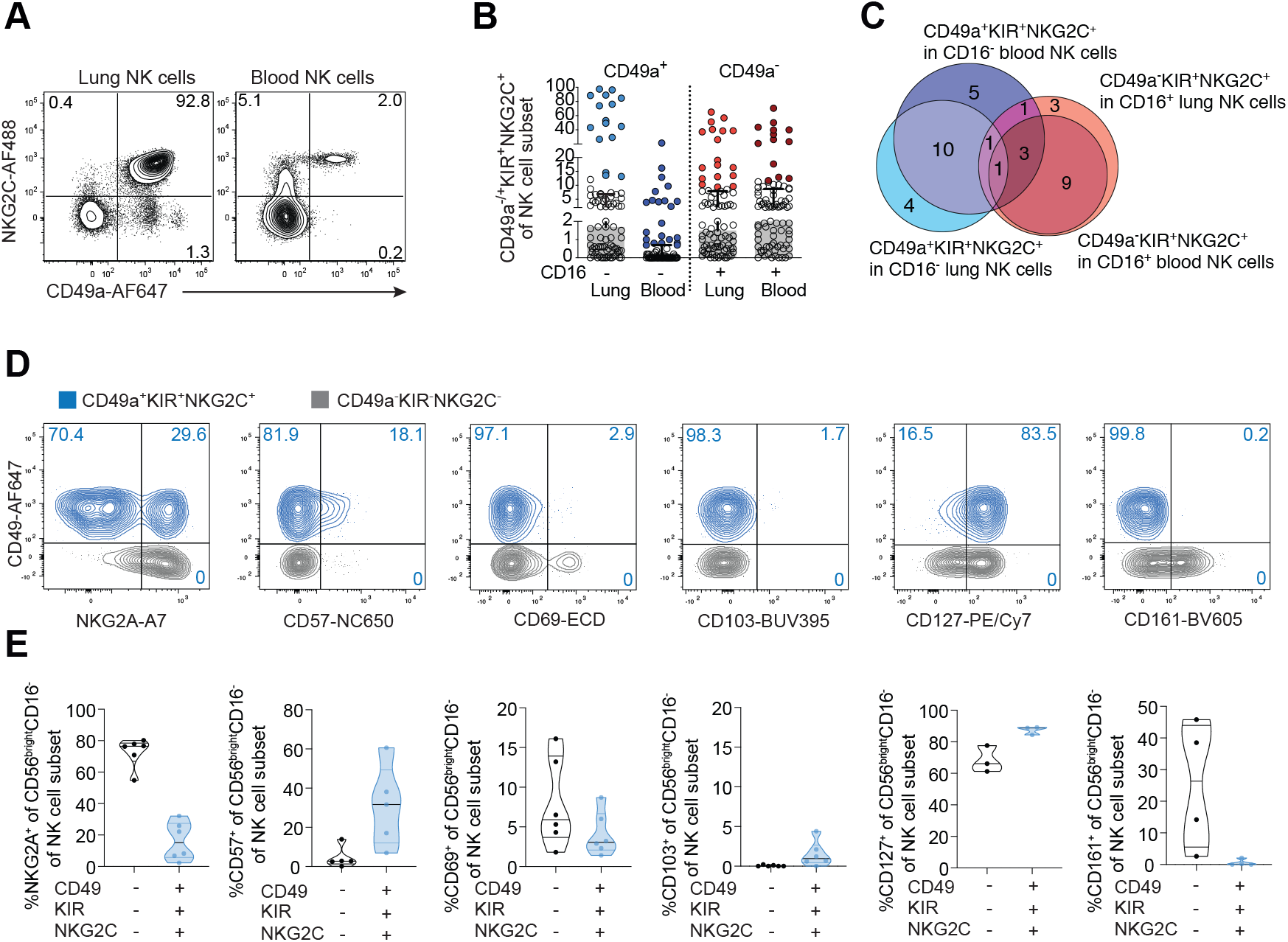
Expansions of adaptive-like trNK cell in the lung indicate presence of adaptive-like CD49a^+^CD56^bright^CD16^−^ NK cells in paired blood. **(A)** Representative dot plots displaying expression of CD49a and NKG2C on NK cells in lung and paired peripheral blood. **(B)** Summary of data of frequencies of adaptive-like CD49a^+^ NK cells of CD16^−^ trNK cells and of adaptive-like pbNK cells in the CD16^+^ NK cell subset in paired lung and peripheral blood. Adaptive-like NK cell “expansions” were identified as outliers (filled circles) using the Robust regression and Outlier removal (ROUT) method (ROUT coefficent Q=1). Error bars show the median with interquartile range (n = 86). Median with interquartile range is shown. **(C)** Euler diagram indicating overlaps and relationships between adaptive-like trNK and pbNK cell expansions in peripheral blood and lung. The number of individuals with overlaps between the subsets and compartments are indicated in the circles. **(D)** Representative overlays and **(E)** summary of data showing phenotypic differences between adaptive-like CD49a^+^ (blue) and non-adaptive CD49a^−^ (grey) NK cells within the CD56^bright^CD16^−^ NK cell subset in blood. (NKG2A, n=6; CD57, n=5; CD69, n=6; CD103, n=6; CD127, n=3; CD161, n=4). Violin plots with quartiles and median are shown.

Taken together, these results demonstrate the presence of an adaptive-like CD49a^+^CD56^bright^CD16^−^ NK cell subset in the peripheral blood of a subset of donors, which is linked to adaptive-like trNK cells in the human lung and emerging independently from CD56^dim^CD16^+^ adaptive-like pbNK cells.

### Peripheral blood adaptive-like CD49a^+^CD56^bright^CD16^−^ pbNK cells from healthy donors share features with lung trNK cells and adaptive-like CD56^dim^CD16^+^ pbNK cells

The presence of adaptive-like lung trNK cells in patients undergoing surgery for suspected lung cancer did not significantly correlate with any demographical or clinical parameters including age, gender, cigarette smoking, COPD, the type of lung tumor, survival, lung function, or HCMV IgG concentrations in plasma (Supplementary Fig. 2A-F). Since we could identify circulating CD49a^+^CD56^bright^CD16^−^ pbNK cells in the majority of patients with an expansion of adaptive-like trNK cells in the lung, we next sought to determine if such cells could also be detected in the peripheral blood of unrelated healthy donors. Indeed, we found KIR^+^NKG2C^+^ NK cells co-expressing CD49a in the CD56^bright^CD16^−^ NK cell subset in 16% of healthy blood donors (Fig. 5A, B). The frequencies of CD49a^+^CD56^bright^CD16^−^ pbNK cells out of CD16^−^ NK cells were lower in healthy peripheral blood (up to 5%, mean 0.3%) as compared to those found in patients with suspected lung cancer (up to 21%, mean 1.2%) (Fig. 5B). Within the CD56^bright^CD16^−^ NK cell subset, KIR^+^NKG2C^+^ NK cells were almost exclusively detected in the CD49a^+^ population (Fig. 5C). UMAP analysis of CD56^bright^CD16^−^ NK cells from healthy donors with CD49a^+^CD56^bright^CD16^−^ NK cells in the blood revealed a strong separation of the CD49a^+^ NK cell subset co-expressing KIR and NKG2C based on lower expression or lack of CD69, CD45RA, CD57, CD38, NKp80, and TIM-3 as well as high expression of CD8, CXCR3 and granzyme B on CD49a^+^KIR^+^NKG2C^+^ NK cells (Fig. 5D). This phenotype could be confirmed in individual samples (Fig. 5E). Interestingly, strong expression of CXCR6 could be identified on CD69^+^, but not on adaptive-like CD49a^+^ CD56^bright^CD16^−^ pbNK cells, indicating that the latter NK cell subset depends on other chemokine receptors such as CXCR3 for tissue homing.

**Figure 5:**
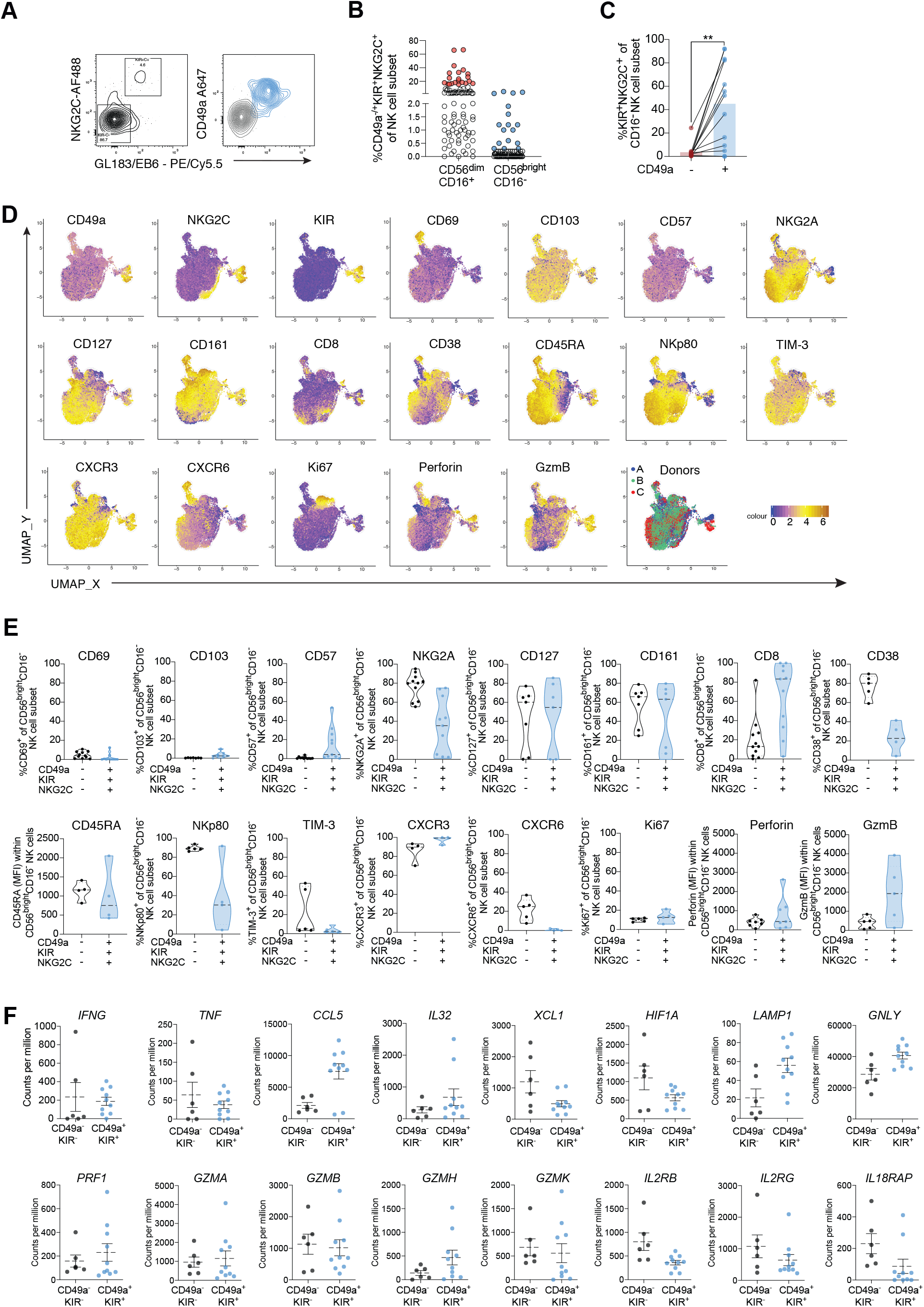
Expansions of adaptive-like CD49a^+^CD56^bright^CD16^−^ NK cells in healthy blood donors. **(A)** Representative dot plot (left plot) and overlay (right plot) showing expression of KIR and NKG2C (left plot), and CD49a on adaptive-like KIR^+^NKG2C^+^ NK cells (blue) versus non-adaptive KIR^−^NKG2C^−^ NK cells (grey) (right plot) within CD16^−^ blood NK cells of healthy blood donors. **(B)** Identification of expansions (filled circles) of adaptive-like CD56^dim^CD16^+^ pbNK cells (21 outliers, 20%) and adaptive-like CD49a^+^CD56^bright^CD16^−^ pbNK cells (17 outliers, 16%) via the ROUT method (see also Figure 4E). Error bars show the median with interquartile range. (n=95). Median with interquartile range is shown. **(C)** Frequencies of KIR^+^NKG2C^+^ cells of CD49a^−^ CD16^−^ or CD49a^+^CD16^−^ NK cells in healthy blood. The respective maternal population comprised at least 45 cells (n=13). Wilcoxon matched-pairs signed rank test. **p<0.005 **(D)** UMAPs based on CD56^bright^CD16^−^ NK cells from three donors with KIR^+^NKG2C^+^CD56^bright^CD16^−^ NK cells. UMAPs were constructed using expression of CXCR3, CD161, Ki67, NKG2C, CD103, TIGIT, perforin, granzyme B, NKG2A, CD16, CD56, CD49a, CD38, CD8, CXCR6, CD4, CD57, CD45RA, NKp80, CD69, GL183/EB6 (KIR), and CD127. Color scale indicates log2(normalized protein expression +1) for each parameter. **(E)** Summary of protein expression on adaptive-like CD49a^+^ NK cells from peripheral blood from healthy donors. (CD69, n=11; CD103, n=7; CD57, n=11; NKG2A, n=11; CD127, n=7; CD161, n=7; CD8, n=11; CD38, n=5; CD45RA, n=4; NKp80, n=5; TIM-3, n=5; CXCR3, n=4; CXCR6, n=5; Ki67, n=5; perforin, n=7; granzyme B, n=5). Violin plots with quartiles and median are shown. **(F)** Gene expression levels (counts per million reads) for selected genes associated with functional capacity are shown for CD49a^−^KIR^−^ and CD49a^+^KIR^+^ blood CD56^bright^CD16^−^ NK cells. Mean ± SEM is shown.

To gain further insight into the adaptive-like CD49a^+^ pbNK cells, we sorted this subset and compared it to sorted blood non-adaptive CD49a^−^CD56^bright^CD16^−^ NK cells using RNAseq (Fig. 6A, see Supplementary Fig. 1C for gating strategy). We next investigated whether gene expression differences in adaptive-like CD49a^+^ pbNK cells indicated a different functional profile. Adaptive-like CD49a^+^ pbNK cells expressed particularly higher levels of *CCL5*, *LAMP1, GZMH* and *GNLY*, and lower levels of *XCL1*, *HIF1A*, *IL2RB*, and *L18RAP* (Fig. 5F). Hence, adaptive-like CD49a^+^ pbNK cells and adaptive-like lung trNK cells from different donors (Fig. 3A) shared a common gene expression pattern for some (*CCL5*, *GZMH*, *IL2RB* and *IL18RAP)*, but not all (i.e., *GNLY*) genes, indicating that they are functionally distinct from each other but also from their non-adaptive counterparts.

**Figure 6:**
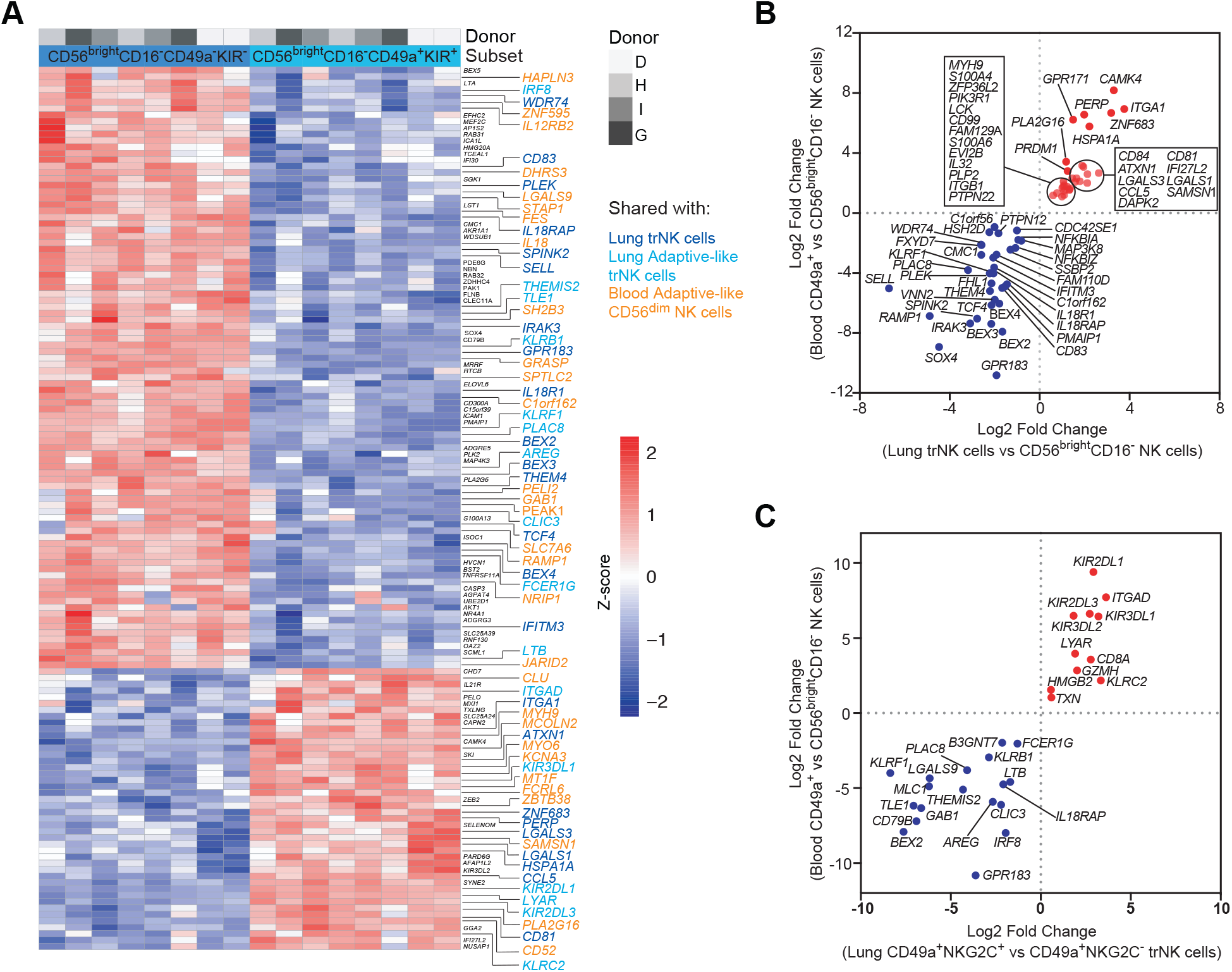
Adaptive-like CD49a^+^CD56^bright^CD16^−^ pbNK cells in healthy blood donors share traits with both trNK cells and with adaptive-like trNK cells. **(A)** Heatmap showing 138 differentially expressed genes (padj<0.001, log2FC>2) between adaptive-like CD49a^+^CD56^bright^CD16^−^ NK cells and non-adaptive CD49a^−^ CD56^bright^CD16^−^ NK cells in peripheral blood from unrelated healthy donors (n=4). Genes shared with trNK cells in the lung were highlighted in dark blue, shared with adaptive-like trNK cells in bright blue, and genes shared with adaptive-like CD56^dim^CD16^+^ pbNK cells in orange. **(B)** Log2 fold-change for trNK cells vs non-tissue-resident CD56^bright^CD16^−^ NK cells in lung against log2 fold-change for adaptive-like CD49a^+^CD56^bright^CD16^−^ vs non-adaptive CD49a^−^CD56^bright^CD16^−^ NK cells in blood. **(C)** Log2 fold-change for KIR^+^NKG2C^+^ trNK cells vs NKG2C^−^ trNK cells in lung against log2 fold-change for adaptive-like CD49a^+^CD56^bright^CD16^−^ vs non-adaptive CD49a^−^CD56^bright^CD16^−^ NK cells in blood.

To assess whether adaptive-like CD49a^+^CD56^bright^CD16^−^ pbNK cells segregate further at the transcriptome level, we analyzed differentially expressed genes between adaptive-like CD49a^+^CD56^bright^CD16^−^ pbNK cells and non-adaptive CD56^bright^CD16^−^ NK cells in peripheral blood from healthy blood donors. A total of 351 genes were differentially expressed (padj<0.01, log2FC>1) and clearly segregated both subsets (Fig. 6A). Since adaptive-like CD49a^+^ pbNK cells resembled to some extent adaptive-like trNK cells in the lung, we next sought to identify similarities to non-adaptive and adaptive-like trNK cells also at transcriptome level. For this, we compared differentially expressed genes in adaptive-like CD49a^+^CD56^bright^CD16^−^ pbNK cells (compared to CD49a^−^KIR^−^CD56^bright^CD16^−^ NK cells in peripheral blood) and in non-adaptive lung trNK cells (defined as CD69^+^CD49a^+^CD103^+^NKG2A^+^NKG2C^−^CD16^−^ NK cells) (compared to non-tissue-resident CD69^−^CD56^bright^CD16^−^ NK cells in lung) (Fig. 6B). Adaptive-like CD49a^+^CD56^bright^CD16^−^ pbNK cells shared 73 DEGs with non-adaptive trNK cells in lung, including high expression of *ITGA1* (CD49a), *ZNF683* (Hobit), *PRDM1* (Blimp-1), *CCL5*, *PIK3R1*, *PLA2G16*, *ATXN1*, as well as lower expression of *SELL* (CD62L), *GPR183*, *IL18R1*, *IL18RAP*, *SOX4*, *RAMP1*, and *IFITM3* (Fig. 6B). All of these genes have also been shown to be differentially expressed in trNK cells in the bone marrow and/or CD8^+^ T_RM_ cells in lung (32,34). It should however be noted that other core-genes associated with tissue-resident lymphocytes (e.g. *S1PR1*, *S1PR5*, *CXCR6*, *ITGAE*, *RGS1*, *KLF2*, *KLF3*, and *RIPOR2*) were not differentially expressed between adaptive-like CD49a^+^CD56^bright^CD16^−^ pbNK cells and non-adaptive CD56^bright^CD16^−^NK cells, indicating that they only partially have a tissue-resident phenotype.

Next, we determined whether adaptive-like CD49a^+^CD56^bright^CD16^−^ pbNK share a common gene signature with adaptive-like trNK cells and/or CD56^dim^CD16^+^ pbNK cells. Indeed, adaptive-like CD49a^+^CD56^bright^CD16^−^ pbNK cells shared differentially expressed genes with adaptive-like trNK cells in lung (Fig. 6C) and/or CD56^dim^CD16^+^ pbNK cells (8,25), including increased expression of *KIRs*, *KLRC2*, *GZMH*, *ITGAD*, *CCL5*, *IL32*, *ZBTB38*, *CD3E*, *ARID5B*, *MCOLN2*, and *CD52,* and decreased expression of *KLRB1*, *FCER1G*, *IL18RAP*, *IL2RB2*, *TLE1*, *AREG*, and *KLRF1* (Fig. 6A, C, Supplementary Fig. 3).

Taken together, adaptive-like CD49a^+^CD56^bright^CD16^−^ pbNK cells share traits with both non-adaptive lung trNK cells, and adaptive-like trNK and CD56^dim^CD16^+^ pbNK cells.

## Discussion

Human adaptive-like NK cells have been described within the CD56^dim^CD16^+^ subset in peripheral blood (5,6,8,10,35) and the CD56^bright^CD16^−^ subset in liver (7). Here, we identified and characterized a yet unexplored and unique subset of adaptive-like CD49a^+^KIR^+^NKG2C^+^CD56^bright^CD16^−^ trNK cells in the human lung, paired blood, and in unrelated healthy human blood. Lung adaptive-like trNK cells shared several phenotypic features with other adaptive-like NK cell subsets both in blood and/or liver, including high expression of CD49a (liver), CD69 (liver), CD2 (blood) and lack of, or decreased expression of CD57 (liver), CD45RA (liver) and perforin (liver), as well as low expression of FcεR1γ and Siglec-7 (blood) (5,7–10,21,35,36). However, lung adaptive-like trNK cells segregate from liver adaptive-like trNK cells on the basis of high expression of Eomes and CD103, and from adaptive-like CD56^dim^CD16^+^ pbNK cells by lack of CD57 and a CD56^bright^CD16^−^ phenotype (7). Transcriptome analysis revealed shared core genes in adaptive-like trNK cells, and CD56^dim^CD16^+^ and CD56^bright^CD16^−^ pbNK cells, underlining common features between all adaptive-like NK cell populations. Intriguingly, lung adaptive-like trNK cells were highly target cell-responsive, and the overall paired blood and lung NK cell populations were hyperresponsive in donors with expansions of adaptive-like trNK cells in the lung. These findings indicate *in vivo* priming akin to what has been observed previously in human antigen-dependent (3,37), antigen-independent, and cytokine-dependent (2,38–40) NK cell recall responses. Furthermore, adaptive-like CD56^dim^CD16^+^ pbNK cells have recently been shown to be functionally primed against target cells following IL-12 and IL-18 stimulation, while showing only a poor response to these cytokines alone (41). In line with our results, this emphasizes a role of potential cytokine-mediated priming of adaptive-like trNK cells in the human lung.

Despite the indications of *in vivo* priming, the presence of expansions of adaptive-like CD49a^+^CD56^bright^CD16^−^ NK cells in the lung donors did not correlate with presence or kind of tumor, HCMV serostatus, or clinical and demographic parameters. In fact, we could identify adaptive-like CD49a^+^CD56^bright^CD16^−^ NK cells also in the peripheral blood of healthy donors. These adaptive-like CD49a^+^CD56^bright^CD16^−^ pbNK cells shared a gene signature with trNK cells in the human lung, indicating tissue imprinting. These findings suggest re-entry of adaptive-like trNK cells from tissue into circulation and, hence, potential seeding of tissues with adaptive-like trNK cells via peripheral blood. In mice, CD8^+^ T effector cells egress from infected lung in a tightly regulated manner following infection with influenza A virus (42), and CD8^+^ T_RM_ cells wane over time in self-limiting viral infections of the respiratory tract (43). MCMV-specific CD8^+^ T cells convertg to CD103^+^ T_RM_ cells, with small numbers of new T_RM_ cells deriving from the circulation (44), and memory inflation is required for retention of CD8^+^ T_RM_ cells in the lungs after intranasal vaccination with MCMV (45). This indicates a dynamic retention of T_RM_ cells by a persistent infection. It remains to be determined whether virus-dependent expansion and maintenance of T_RM_ cells is analogous in adaptive-like trNK cells in the lung. However, our data indicate that T_RM_ cells and adaptive-like NK cells differ at least in their recruitment to the lung, with T_RM_ cells being dependent on CXCR6 (46) while adaptive-like CD49a^+^CD56^bright^CD16^−^ pbNK cells lacked CXCR6 but expressed high levels of CXCR3.

Adaptive-like CD49a^+^CD56^bright^CD16^−^ NK cell expansions were rarely observed in donors with adaptive-like CD56^dim^CD16^+^ pbNK cell expansions, indicating that these two distinct subsets have different developmental cues. Indeed, even in the rare cases where we could detect expansions of both adaptive-like CD49a^+^CD56^bright^CD16^−^ NK cells and CD56^dim^CD16^+^ pbNK cells in the same individual, these populations displayed specific and individual KIR repertoires. Furthermore, expansions of adaptive-like lung trNK cells were detected in HCMV-seronegative individuals (Supplementary Fig. 2E, F), while expansions of adaptive-like CD56^dim^CD16^+^ pbNK cells were restricted to HCMV-seropositive individuals (Supplementary Fig. 2E) (5,35). It thus remains possible that other viral infections than CMV in humans and mice (1,47,48) could drive the expansion of adaptive-like trNK cells, as has previously been suggested for the generation of cytokine-induced memory NK cells, e.g. after influenza virus infection in humans (49) and mice (4,50,51), as well as vesicular stomatitis virus (VSV) (4), vaccinia virus (52), HIV-1 (4), and herpes simplex virus 2 (HSV-2) (51), and after immunization with simian immunodeficiency virus (SIV) in rhesus macaques (53). Taken together, our data support a model where adaptive-like trNK cells and CD56^dim^CD16^+^ pbNK cells develop independently from each other, possibly due to distinct environmental requirements for their expansion.

We observed increased gene expression levels of *GZMH* in adaptive-like CD49a^+^CD56^bright^CD16^−^ lung and blood NK cells, and levels for *CCL5* were higher in both conventional and adaptive-like lung trNK cells as compared to CD69^−^ CD56^bright^CD16^−^ lung NK cells, and particularly highly expressed in adaptive-like CD49a^+^CD56^bright^CD16^−^ pbNK cells. An antiviral activity has been proposed for granzyme H (54,55), however, a direct association of this effector molecule with adaptive-like NK cells remains to be determined. In contrast, *CCL5* and *XCL1*, which are both upregulated in human CD49a^+^CD56^bright^CD16^−^ adaptive-like NK cells, were predominantly produced by mouse Ly49H^+^ NK cells upon stimulation with MCMV-derived m157 protein (56), and CCL5 has been shown to be specifically expressed by CD8^+^ memory T_EM_ cells (57). Thus, adaptive-like CD49a^+^CD56^bright^CD16^−^ blood and lung NK cells share functional characteristics with other memory lymphocyte populations. Furthermore, we showed that adaptive-like lung trNK cells were hyperresponsive against target cells, hence, they might be clinically relevant e.g. in disease progression in respiratory viral infections and/or the defense against malignant tumor cells. Similarly, lung CD8^+^ T_RM_ cells have previously been shown to be able to control tumor growth and to correlate with increased survival in lung cancer patients (58). Since adaptive-like trNK cells likely exceed their circulating pbNK cell counterpart in tissue-homing and tumor-infiltration based on their expression of tissue-specific receptors such as CD49a and CXCR3, these cells could be harnessed for future treatment options of solid tumors.

Together, our data reveal the presence of a yet unexplored and distinct adaptive-like trNK cell subset in the human lung, indicating that adaptive-like NK cells are not confined to peripheral blood and/or liver and that different lineages of adaptive-like NK cells potentially exist. Expansions of adaptive-like trNK cells in the lung were commonly accompanied by the presence of adaptive-like CD49a^+^CD56^bright^CD16^−^ NK cells in paired peripheral blood, enabling the non-invasive identification of donors with potential adaptive-like lung trNK cell expansions as well as the isolation of adaptive-like NK cells with tissue-resident characteristics. Finally, adaptive-like NK cells with tissue-resident features and excessive functional responsiveness in the human lung and blood could be an attractive source for tailored cancer immunotherapies, in particular for targeting solid tumors.

## Abbreviations

Eomes: Eomesodermin
FITC: Fluorescein isothiocyanate
HCMV: Human cytomegalovirus
ILC: Innate lymphoid cell
KIR: Killer cell immunoglobulin-like receptor
NK: Natural killer
PE: Phycoerythrin
ROS: Reactive oxygen species

## Acknowledgements

The authors acknowledge support from the National Genomics Infrastructure in Stockholm funded by Science for Life Laboratory, the Knut and Alice Wallenberg Foundation, the Swedish Research Council, and SNIC/Uppsala Multidisciplinary Center for Advanced Computational Science for assistance with massively parallel sequencing and access to the UPPMAX computational infrastructure. The authors also acknowledge the MedH Core Flow Cytometry Facility (Karolinska Institutet), supported by Karolinska Institutet and Region Stockholm, for providing cell-sorting services. Furthermore, the authors want to thank all donors for participating in the study and A.-C. Orre, V. Jackson, and S. Hylander for administrative and clinical help as well as E. Yilmaz for assistance in the laboratory. **Funding:** This work was supported by the Swedish Research Council, the Strategic Research Foundation, the Swedish Foundation for Strategic Research, the Swedish Cancer Society, Sweden’s Innovation Agency, the Eva and Oscar Ahréns Research Foundation, Stockholm, the Åke Wiberg Foundation and the Tornspiran Foundation.

## Author contribution

Conceptualization: N.M., H.G.L., Ja.M.; Methodology: N.M., Ja.M.; Investigation: N.M., M.S., Je.M., J.H., E.K., M.B., S.N., J.N.W.; Resources: M.A.-A.; Writing – original draft: N.M., Ja.M.; Writing – review and editing: N.M., Ja.M., H.G.L; Visualization: N.M., Ja.M.; Funding acquisition: N.M., Ja.M., H.G.L.

## Material and methods

### Lung patients and healthy blood

A total of 103 patients undergoing lobectomy for suspected lung cancer were included in this study for collection of lung tissue and paired peripheral blood. None of the patients received preoperative chemotherapy and/or radiotherapy. Patients with records of strong immunosuppressive medication and/or hematological malignancy were excluded from the study. Clinical and demographic details are summarized in Table 1. Furthermore, healthy blood was collected from regular, unrelated blood donors. The regional review board in Stockholm approved the study, and all donors gave informed written consent prior to collection of samples.

**Table 1.** Clinical and demographic details of the 103 patients included in the study.

|  | **Never-smoker**<br>(n = 16) | **Current smoker**<br>(n = 25) | **Ex-smoker**<br>(n = 62) |
| --- | --- | --- | --- |
| **Female / male** | 8 / 8 | 14 / 11 | 41 / 21 |
| **Age (y), mean ± SD** | 67 ± 13.4 | 67 ± 8.0 | 69 ± 7.1 |
| **FEV1/FVC (% of predicted) mean ± SD** | 100 ± 10.3 | 88 ± 13.7* | 93 ± 14.9<sup>#</sup> |
| **Pathology** | **% (n)** | **% (n)** | **% (n)** |
| Non-malignant | 6 (1) | 4 (1) | 2 (1) |
| Adenocarcinoma | 44 (7) | 52 (13) | 76 (47) |
| Large cell carcinoma | 0 (0) | 12 (3) | 3 (2) |
| Squamous cell carcinoma | 6 (1) | 16 (4) | 5 (3) |
| Metastasis | 0 (0) | 4 (1) | 3 (2) |
| Carcinoid | 31 (5) | 8 (2) | 6 (4) |
| Adenosquamous carcinoma | 6 (1) | 0 (0) | 3 (2) |
| Other<sup>§</sup> | 6 (1) | 4 (1) | 2 (1) |
| **Medication** | **% (n)** | **% (n)** | **% (n)** |
| Inhaled corticosteroids | 19 (3) | 0 (0) | 2 (1) |
| Statins | 38 (6) | 28 (7) | 31 (19) |
| Systemic immunosuppression | 0 (0) | 4 (1) | 5 (3) |
| Beta-agonists or anti-cholinergics | 0 (0) | 12 (3) | 13 (8) |
| Inhaled corticosteroid and long-acting beta-agonist combination | 0 (0) | 8 (2) | 6 (4) |
| **Diagnoses affecting lung function** | **% (n)** | **% (n)** | **% (n)** |
| Asthma | 19 (3) | 0 (0) | 5 (3) |
| COPD | 0 (0) | 32 (8) | 10 (6) |
| Other lung parenchyme disease | 6 (1) | 0 (0) | 2 (1) |

\* n = 24
<sup>#</sup> n = 59
<sup>§</sup> = uncertain histopathological diagnosis, data missing, combined small cell carcinoma

### Processing of tissue specimens and peripheral blood

Lung tissue was processed as previously described (18). Briefly, a small part of macroscopically tumor-free human lung tissue from each patient was transferred into ice-cold Krebs-Henseleit buffer and stored on ice for less than 18 h until further processing. The tissue was digested using collagenase II (0.25 mg/ml, Sigma-Aldrich) and DNase (0.2 mg/ml, Roche), filtered and washed in complete RPMI 1640 medium (Thermo Scientific) supplemented with 10% FCS (Thermo Scientific), 1 mM L-glutamine (Invitrogen), 100 U/ml penicillin, and 50 μg/ml streptomycin (R10 medium). Finally, mononuclear cells from the lung cell suspensions and peripheral blood were isolated by density gradient centrifugation (Lymphoprep).

### RNA-sequencing and RNAseq data analysis

RNA of sorted NK cell subsets from blood and lung were sequenced and analyzed as described previously (15). Briefly, RNAseq was performed using a modified version of the SMART-Seq2 protocol (59). For analysis of lung adaptive-like NK cells, live NKG2C^+^KIR^+^CD3^−^CD14^−^CD19^−^CD56^+^CD16^−^ NK cells were sorted from two donors and were compared to previously published data on CD69^+^CD49a^+^CD103^−^ and CD69^+^CD49a^+^CD103^+^ NKG2A^+^CD16^−^ trNK cells (GSE130379) (15). For analysis of KIR^+^CD49a^+^ CD56^bright^CD16^−^ NK cells, we sorted KIR^+^CD49a^+^ and KIR^−^CD49a^−^ live CD14^−^CD19^−^CD3^−^CD56^bright^CD16^−^ NK cells from cryopreserved PBMCs from 4 donors. Duplicates of 100 cells from each population from two individual donors were sorted into 4.2ul of lysis buffer (0.2% Triton X-100, 2.5uM oligo-dT (5′-AAGCAGTGGTATCAACGCAGAGTACT30VN-3′), 2.5mM dNTP, RNAse Inhibitor (Takara), and ERCC RNA spike in controls (Ambion)) in a 96-well V-bottom PCR plate (Thermo Fisher). Sorted cells were then frozen and stored at −80°C until they could be processed. Subsequent steps were performed following the standard SMART-Seq2 protocol with 22 cycles of cDNA amplification and sample quality was determined using a bioanalyzer (Agilent, High Sensitivity DNA chip). 5ng of amplified cDNA was taken for tagmentation using a customized in-house protocol (60) and Nextera XT primers. Pooled samples were sequenced on a HiSeq2500 on high output mode with paired 2×125bp reads.

### Transcriptome analysis

Following sequencing and demultiplexing, read pairs were trimmed from Illumina adapters using cutadapt (version 1.14) (61), and UrQt was used to trim all bases with a phred quality score below 20 (62). Read pairs were subsequently aligned to the protein coding sequences of the human transcriptome (gencode.v26.pc_transcripts.fa) using Salmon (version 0.8.2) (63), and gene annotation using gencode.v26.annotation.gtf. DeSeq2 (64) was used to analyze RNA-seq data in R studio version 1.20. Briefly, raw count values were used as input into deSeq2, and variance stabilizing transformation was used to transform data. Data were batch- and patient corrected using Limma (65).

A cut-off of >100 counts across the samples was used to filter out low expressed genes. Genes with an adjusted p-value<0.05 and a log2-fold change greater than 1 were considered as differentially expressed between paired samples. Similarly, previously published data sets on adaptive-like NKG2C^+^CD57^+^CD56^dim^CD16^+^ NK cells and conventional NKG2C^−^CD57^+^CD56^dim^ NK cells (GSE117614) (25) were analyzed using deSeq2 to identify differentially expressed genes. Heatmaps of gene expression were generated using Pheatmap in R and show the z-score for differentially expressed genes (as determined above in deSeq2) for all donors and replicates.

### Flow cytometry

Antibodies and clones reactive against the following proteins were used: CD2 (TS1/8, BV421 or Pacific Blue, Biolegend), CD3 (UCHT1, PE-Cy5, Beckman Coulter), CD8 (RPA-T8, Brilliant Violet 570, Biolegend, or RPA-8, BUV395 or SK1, BUV737, BD Biosciences), CD14 (MφP9, Horizon V500, BD Biosciences), CD16 (3G8, Brilliant Violet 570 or Brilliant Violet 711, or Brilliant Violet 785, Biolegend), CD19 (HIB19, Horizon V500, BD Biosciences), CD38 (HIT2, Brilliant Violet 711 or BUV661, BD Biosciences), CD45 (HI30, Alexa Fluor 700, Biolegend, or BUV805, BD Biosciences), CD45RA (HI100, Brilliant Violet 785, Biolegend), CD49a (TS2A, AlexaFluor 647, Biolegend, or HI30, BUV615, or 8R84, Brilliant Violet 421, BD Biosciences), CD56 (N901, ECD, Beckman Coulter, or HCD56, Brilliant Violet 711, Biolegend, or NCAM16.2, PE-Cy7, or BUV563, BD Biosciences), CD57 (TB01, purified, eBioscience, or HNK-1, Brilliant Violet 605, Biolegend), CD103 (APC, B-Ly7, eBioscience, or biotin, 2G5, Beckman Coulter, or Ber-ACT8, Brilliant Violet 711, BUV395, BD Biosciences, or Ber-ACT8, PE-Cy-7, Biolegend), KIR2DL1 (FAB1844F, biotin, R&D Systems), KIR2DL3 (180701, FITC, R&D Systems), KIR3DL1 (DX9, Brilliant Violet 421, Biolegend), KIR3DL2 (DX-31, Brilliant Violet 711, Biolegend), KIR2DL2/S2/L3 (GL183, PE-Cy5.5, Beckman Coulter), KIR2DL1/S1 (EB6, PE-Cy5.5 or PE-Cy7, Beckman Coulter), NKG2A (Z1991.10, APC-A780, or PE, Beckman Coulter, or 131411, BUV395, BD Biosciences), NKG2C (134591, Alexa-Fluor 488 or PE, R&D Systems), CD69 (TP1.55.3, ECD, Beckman Coulter, or FN50, PE-CF594, BD Biosciences, or FN50, Brilliant Violet 786, Biolegend), CD127 (Brilliant Violet 421, HIL-7R-M21, BD Biosciences or PE-Cy7, R34.34, Beckman Coulter), CD161 (HP3-3G10, Brilliant Violet 605 or APC/Fire 750, Biolegend), CXCR3 (Alexa Fluor 647, G025H7, Biolegend), CXCR6 (K041E5, Brilliant Violet 421, Biolegend), CD85j/ILT2 (HP-F1, Super Bright 436, Invitrogen), NKp80 (5D12, PE, BD Biosciences, or 4A4.D10, PE-Vio770, Miltenyi), Siglec-7 (5-386, Alexa Fluor 488, Bio-Rad), TIM-3 (7D3, Brilliant Violet 711, BD Biosciences). After two washes, cells were stained with streptavidin Qdot 605 or Qdot 585 (both Invitrogen), anti-mouse IgM (II/41, eFluor 650NC, eBioscience) and Live/Dead Aqua (Invitrogen). After surface staining, peripheral blood mononuclear cells (PBMC) were fixed and permeabilized using FoxP3/Transcription Factor staining kit (eBioscience). For intracellular staining the following antibodies were used: Eomes (WF1928, FITC, eBioscience), FcεR1γ (polyclonal, FITC, Merck), granzyme B (GB11, BB790, BD Biosciences), Ki67 (B56, Alexa Fluor 700, BD Biosciences), perforin (dG9, BB755, BD Biosciences, or B-D48, Brilliant Violet 421, Biolegend), T-bet (4B10, Brilliant Violet 421, BD Biosciences), and TNF (MAb11, Brilliant Violet 421, Biolegend, or Brilliant Violet 650, BD Biosciences). Purified NKG2C (134591, R&D Systems) was biotinylated using a Fluoreporter Mini-biotin XX protein labeling kit (Life Technologies) and detected using streptavidin-Qdot 585, 605 or 655 (Invitrogen).

Samples were analyzed on a BD LSR Fortessa equipped with four lasers (BD Biosciences) or a BD FACSymphony A5 equipped with five lasers (BD Biosciences), and data were analyzed using FlowJo version 9.5.2 and version 10.6.1 (Tree Star Inc). UMAPs were constructed in FlowJo 10.6.1 using the UMAP plugin. UMAP coordinates and protein expression data were subsequently exported from FlowJo, and protein expression for each parameter was normalized to a value between 0 and 100. UMAP plots were made in R using ggplot, and color scale show log2(normalized protein expression +1).

For sorting of NK cells from lung and peripheral blood for RNA sequencing, thawed cryopreserved mononuclear cells were stained with anti-human CD57 (NK-1, FITC, BD Biosciences), CD16 (3G2, Pacific Blue, BD Biosciences), CD14 (MφP9, Horizon V500, BD Biosciences), CD19 (HIB19, Horizon V500, BD Biosciences), CD103 (Ber-ACT8, Brilliant Violet 711, BD Biosciences), CD49a (TS2/7, Alexa Fluor 647, Biolegend), CD45 (HI30, A700, Biolegend), CD8 (RPA-T8, APC/Cy7, BD Biosciences), NKG2A (Z199.10, PE, Beckman Coulter), CD69 (TP1.55.3, ECD, Beckman Coulter), CD3 (UCHT1, PE/Cy5, Beckman Coulter), KIR2DL1/S1 (EB6, PE/Cy5.5, Beckman Coulter), KIR2DL2/3/S2 (GL183, PE/Cy5.5, Beckman Coulter,), NKG2C (134591, biotin, R&D Systems, custom conjugate), CD56 (NCAM16.1, PE/Cy7, BD Biosciences), streptavidin Qdot655 (Invitrogen), and Live/Dead Aqua (Invitrogen).

### DNA isolation and KIR/HLA-ligand genotyping

Genomic DNA was isolated using a DNeasy Blood & Tissue Kit (Qiagen) from 100 μl of whole blood. KIR genotyping and KIR ligand-determination were performed using PCR-SSP technology with a *KIR* typing kit and a *KIR HLA* ligand kit (both Olerup-SSP) according to the manufacturer’s instructions.

### CMV IgG ELISA

Concentrations of anti-CMV IgG relative to a standard curve and internal negative and positive control were determined by ELISA (Abcam, UK) and read in a microplate spectrophotometer (Bio-Rad xMark) at 450nm with a 620nm reference wavelength.

### Activation assay

Degranulation and TNF production of fresh blood and lung NK cells were assessed as previously described (18,19). In brief, fresh lung and blood mononuclear cells were resuspended in R10 medium and rested for 15 to 18 hours at 37°C. Subsequently, the cells were co-cultured in R10 medium alone or in presence of K562 cells for 2 hours in the presence of anti-human CD107a (FITC or Brilliant Violet 421, H4A3, BD Biosciences, San Jose, Calif.).

### Statistics

GraphPad Prism 6 and 7 (GraphPad Software) was used for statistical analysis. For each analysis, measurements were taken from distinct samples. The statistical method used is indicated in each figure legend.

#### Data Availability

The dataset generated for this study can be found in the Gene Expression Omnibus with accession no. xxxx (data will be deposited and made available before publication).

