## Supplemental Fig. 1 for "Expansions of adaptive-like NK cells with a tissue-resident phenotype in human lung and blood"

Supplementary Figure 1

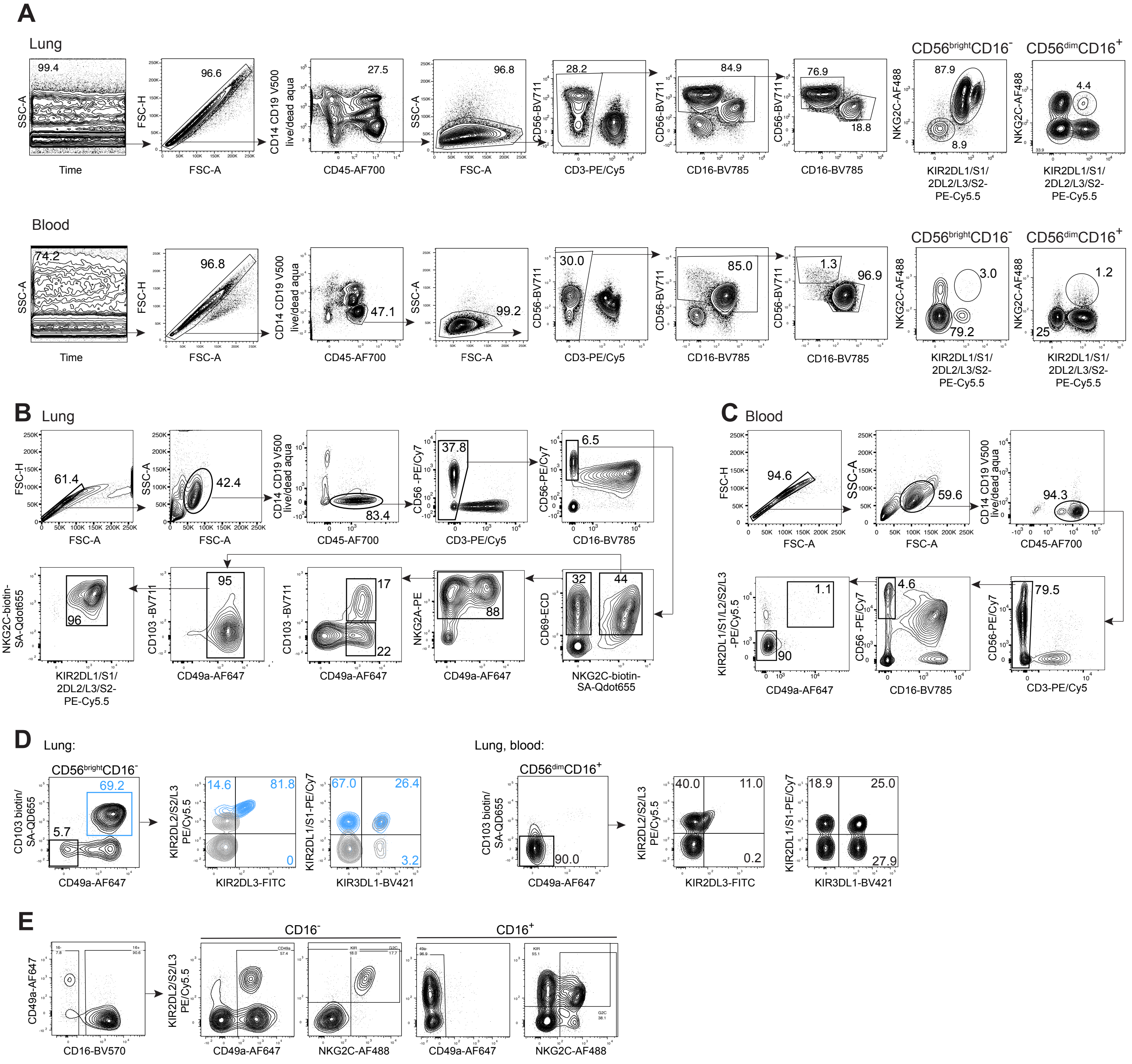


**Supplementary Figure 1: Identification of adaptive-like NK cell subsets in human lung and peripheral blood.** **(A)** Gating strategy to identify KIR$^{+}$NKG2C$^{+}$ NK cells in human lung (upper panel) and blood (lower panel) (related to Fig. 1, 2A-D, 3B). **(B)** Gating strategy for sort for RNA sequencing analysis of lung (related to Fig. 2E,F, 3A) and **(C)** healthy blood NK cells (related to Fig. 5F, 6). **(D)** Gating strategy for single KIR analysis on CD56$^{bright}$CD16$^{-}$ lung NK cells (left panel) and CD56$^{dim}$CD16$^{+}$ NK cells in lung and blood (right panel) (related to Fig. 2G-I). **(E)** Gating strategy for identification of outliers within the CD16$^{-}$ and CD16$^{+}$ NK cell subsets (related to Fig. 4B,C).
