## Supplemental Fig. 2 for "Expansions of adaptive-like NK cells with a tissue-resident phenotype in human lung and blood"

Supplementary Figure 2

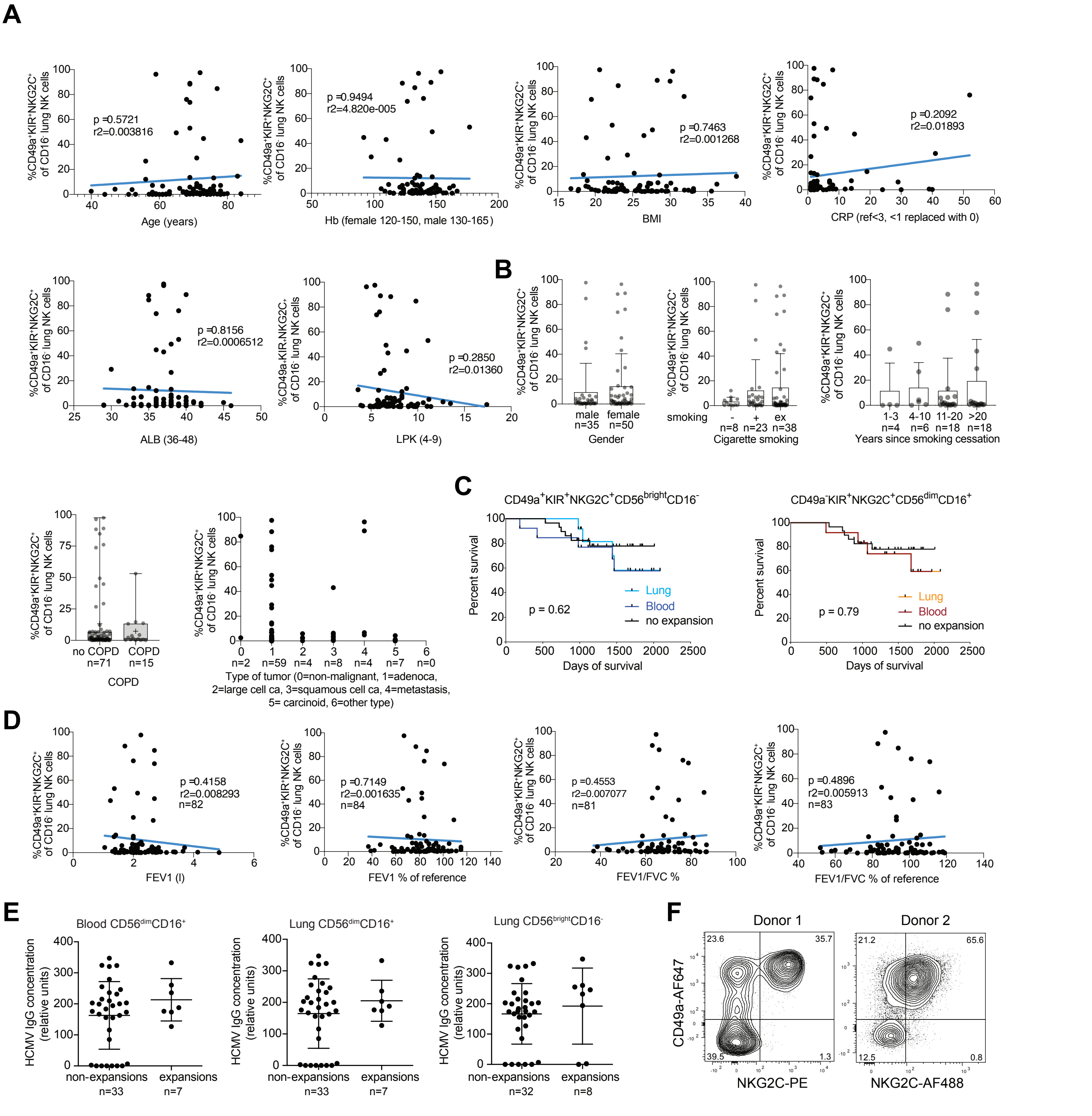


**Supplementary figure 2**: Association of adaptive trNK cells and patient characteristics. **(A)** Linear regression analysis of the frequency of CD49a$^{+}$KIR$^{+}$NKG2C$^{+}$ cells among CD16$^{-}$ lung NK cells versus age, hemoglobin (Hb) levels, body mass index (BMI), c-reactive protein (CRP) levels, albumin (ALB) levels, and leucocyte count (LPK) (n =86). **(B)** Frequencies of CD49a$^{+}$KIR$^{+}$NKG2C$^{+}$NK cells among CD16$^{-}$ lung NK cells versus gender, cigarette smoking status, years since cigarette smoking cessation, chronic obstructive pulmonary disease (COPD) status, and type of tumor. Mean +/- SD is shown. **(C)** Days of survival in donors with CD49a$^{+}$KIR$^{+}$NKG2C$^{+}$CD56$^{bright}$CD16$^{-}$ NK cell expansions in blood and/or lung (left plot) and in donors with CD49a$^{+}$KIR$^{+}$NKG2C$^{+}$CD56$^{bright}$CD16$^{-}$ NK cell expansions in blood and/or lung (right plot) (n=57). **(D)** Linear regression analysis of FEV1 (l), FEV1 % of reference, FEV1/FV%, and FEV1/FVC % of reference is shown. **(E)** HCMV IgG concentration in plasma from donors with or without KIR$^{+}$NKG2C$^{+}$CD56$^{dim}$CD16$^{+}$ expansions in blood (left) and lung (middle) and in donors with KIR$^{+}$NKG2C$^{+}$CD56$^{dim}$CD16$^{+}$ expansions in lung (right). **(F)** Expression of NKG2C and CD49a on CD56$^{bright}$CD16$^{-}$ NK cells from the two HCMV-seronegative donors in (e) demonstrate adaptive-like expansions of CD56$^{bright}$CD16$^{-}$ NK cells in the absence of HCMV seroconversion.
