## Supplemental Fig. 3 for "Expansions of adaptive-like NK cells with a tissue-resident phenotype in human lung and blood"

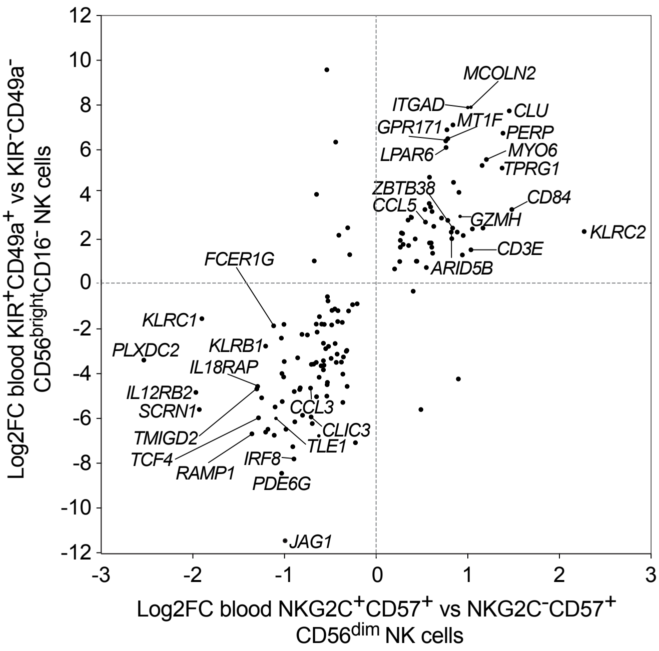


**Supplementary Figure 3: Comparison of DEGs between blood CD49a⁺KIR⁺ CD56^bright^CD16⁻ NK cells and CD57⁺NKG2C⁺CD56^dim^ NK cells.** Log2-fold changes in gene expression for shared significantly differentially expressed genes between blood $CD49a^{+}KIR^{+}$ and $CD49a^{-}KIR^{-}$ $CD56^{bright}CD16^{-}$ NK cells (y-axis) and $CD57^{+}NKG2C^{+}$ and $CD57^{+}NKG2C^{-}$ $CD56^{dim}$ NK cells (x-axis). Data for $CD56^{dim}$ NK cells are from GSE117614 (Cichocki et al).
